# Few-shot learning for highly accelerated 3D time-of-flight MRA reconstruction

**DOI:** 10.1101/2025.04.14.648710

**Authors:** Hao Li, Mark Chiew, Iulius Dragonu, Peter Jezzard, Thomas W. Okell

**Author notes:** Correspondence: Thomas W. Okell, Wellcome Centre for Integrative Neuroimaging, FMRIB Division, Nuffield Department of Clinical Neurosciences, John Radcliffe Hospital, Headley Way, Headington, Oxford, OX3 9DU, UK. **Funding information** Wellcome Trust, Grant/Award Numbers: 203139/Z/16/Z, 203139/A/16/Z, 220204/Z/20/Z; NIHR Oxford Biomedical Research Centre, Grant/Award Number: NIHR203311; NIHR Oxford Health Biomedical Research Centre, Grant/Award Number NIHR203316; Siemens Healthineers; Medical Research Council; Canada Research Chairs; Dunhill Medical Trust.

## Abstract

**Purpose:** To develop a deep learning-based reconstruction method for highly accelerated 3D time-of-flight magnetic resonance angiography (TOF-MRA) that achieves high-quality reconstruction with robust generalization using extremely limited acquired raw data, addressing the challenge of time-consuming acquisition of high-resolution, whole-head angiograms.

**Methods:** A novel few-shot learning-based reconstruction framework is proposed, featuring a 3D variational network specifically designed for 3D TOF-MRA that is pre-trained on simulated complex-valued, multi-coil raw k-space datasets synthesized from diverse open-source magnitude images and fine-tuned using only two single-slab experimentally acquired datasets. The proposed approach was evaluated against existing methods on acquired retrospectively undersampled *in vivo* k-space data from five healthy volunteers and on prospectively undersampled data from two additional subjects.

**Results:** The proposed method achieved superior reconstruction performance on experimentally acquired *in vivo* data over comparison methods, preserving most fine vessels with minimal artifacts with up to 8-fold acceleration. Compared to other simulation techniques, the proposed method generated more realistic raw k-space data for 3D TOF-MRA. Consistently high-quality reconstructions were also observed on prospectively undersampled data.

**Conclusions:** By leveraging few-shot learning, the proposed method enabled highly accelerated 3D TOF-MRA relying on minimal experimentally acquired data, achieving promising results on both retrospective and prospective *in vivo* data while outperforming existing methods. Given the challenges of acquiring and sharing large raw k-space datasets, this holds significant promise for advancing research and clinical applications in high-resolution, whole-head 3D TOF-MRA imaging.

## 1 Introduction

Angiography, the imaging of blood vessels, is particularly vital in the brain, where ischemia or hemorrhage can have severe clinical consequences^1^. Cerebral angiograms provide critical diagnosis and treatment planning information for various cerebrovascular diseases, including stroke^2^, steno-occlusive disease^3^, and arteriovenous malformation^4^. However, conventional angiographic techniques, such as digital subtraction angiography, rely on X-ray imaging and require contrast agent injection, exposing patients to ionizing radiation and limiting their use in pediatric or longitudinal examinations^5^. Although contrast-enhanced magnetic resonance angiography (CE-MRA) offers a radiation-free alternative, concerns have been raised regarding the accumulation of gadolinium-based contrast agents^6^.

Time-of-flight (TOF) MRA is a non-contrast technique that exploits the magnetization differences between stationary tissue and inflowing blood^7^. With multiple overlapping thin-slab acquisitions^8^, 3D TOF-MRA provides excellent visualization of the arterial vasculature and is routinely used in clinical assessments of cerebrovascular diseases^7^. High-resolution, whole-head 3D TOF-MRA could enable more comprehensive cerebrovascular assessments, improving diagnostic accuracy for small aneurysms^9,10^ and cerebral small vessel diseases^11^. However, acquiring high-resolution whole-brain 3D TOF-MRA requires long scan times, which pose challenges in clinical workflows. Prolonged acquisitions also increase patient discomfort and susceptibility to motion-related artifacts, potentially compromising image quality and diagnostic reliability^12^.

Parallel imaging (PI) techniques, such as GRAPPA^13^ and SENSE^14^, are commonly employed to reduce scan times by leveraging the spatial sensitivity of multiple receiver coils to reconstruct undersampled k-space data. However, PI reconstructions typically degrade at acceleration factors beyond 3–4 due to noise amplification and residual artifacts. Compressed sensing (CS) has been explored extensively for 3D TOF-MRA^15–17^, utilizing incoherent undersampling and sparsity constraints to achieve higher acceleration factors^18^. However, CS reconstructions require lengthy processing times with careful parameter tuning, limiting their clinical application^19^. More recently, wave-controlled aliasing in parallel imaging (wave-CAIPI)^20^ has been applied to 3D TOF-MRA^21^, demonstrating improved image quality at higher acceleration factors compared to conventional PI and CS. However, wave-CAIPI necessitates complex implementation and careful optimization of wave gradient parameters to minimize flow-related artifacts^21^.

Deep learning (DL)-based reconstruction methods have emerged as powerful alternatives for accelerating magnetic resonance imaging (MRI), consistently outperforming CS while significantly reducing reconstruction times^22–24^. A particularly high-performing DL method is the end-to-end variational network (E2E-VarNet)^25^, which incorporates an MRI forward model and uses 2D U-Nets^26^ to learn gradients in unrolled optimization steps. However, these DL-based approaches are primarily designed for and trained on 2D k-space data, which may not be optimal for 3D TOF-MRA, where vessel signals are inherently connected across adjacent slices. Additionally, training such models typically requires large raw k-space datasets, which are generally unavailable for 3D TOF-MRA. Consequently, most previous studies have relied on small in-house datasets^27–30^.

For reconstructing undersampled multi-coil 3D TOF-MRA data, Jun et al. proposed a 3D multi-stream convolutional neural network (CNN) called DPI-Net, which reconstructs directly in the image domain^27^. Chung et al. developed a multiplanar 2D OT-cycleGAN that enables training with unmatched reference data, achieving reconstruction results on an in-house dataset that were comparable to supervised learning approaches trained on paired raw data^28^. Most recently, Sun et al. introduced an uncertainty-aware reconstruction model for accelerated 7T TOF-MRA, utilizing evidential deep learning to support uncertainty quantification^31^.

Data limitations pose significant challenges for effectively training and validating these DL models, which may fail to generalize to different populations, anatomical variations, protocol settings and scanners. Small datasets also increase the risk of overfitting, reducing robustness and reproducibility across institutions^24^. While self-supervised learning has been proposed to address data limitations by leveraging undersampled data, it still commonly requires large undersampled k-space datasets^32–34^. Emerging zero-shot self-supervised methods^35^ attempt to bypass this requirement, but they involve lengthy per-reconstruction training times, making them impractical for large-scale clinical deployment.

To overcome these challenges, this work proposes a DL-based reconstruction framework for highly accelerated 3D TOF-MRA that integrates few-shot learning to eliminate the need for significant numbers of *in-vivo* raw k-space datasets for training. The proposed approach first pre-trains a 3D model, adapted from E2E-VarNet, using simulated raw k-space data generated from diverse publicly available magnitude images. Fine-tuning is then performed using only two experimentally acquired fully sampled *in vivo* single-slab datasets. The proposed method was evaluated against established methods on retrospectively undersampled *in vivo* multi-slab k-space acquisitions from healthy subjects scanned at 3T. Additionally, the effectiveness of the proposed simulation pipeline was assessed in comparison to other raw k-space data synthesizing techniques. Finally, additional prospectively undersampled acquisitions were used to assess the proposed method’s robustness in a real-world clinical scenario. This work builds upon an accepted abstract for presentation at the upcoming 2025 ISMRM Annual Meeting^36^.

## 2 Methods

To effectively train a DL model capable of reconstructing complex-valued multi-coil raw k-space, phase and coil sensitivity maps were simulated to leverage magnitude 3D TOF-MRA images from the large open-source multi-center multi-vendor IXI dataset (https://brain-development.org/ixi-dataset/). An overview of this approach is depicted in Figure 1.

**Figure 1.**
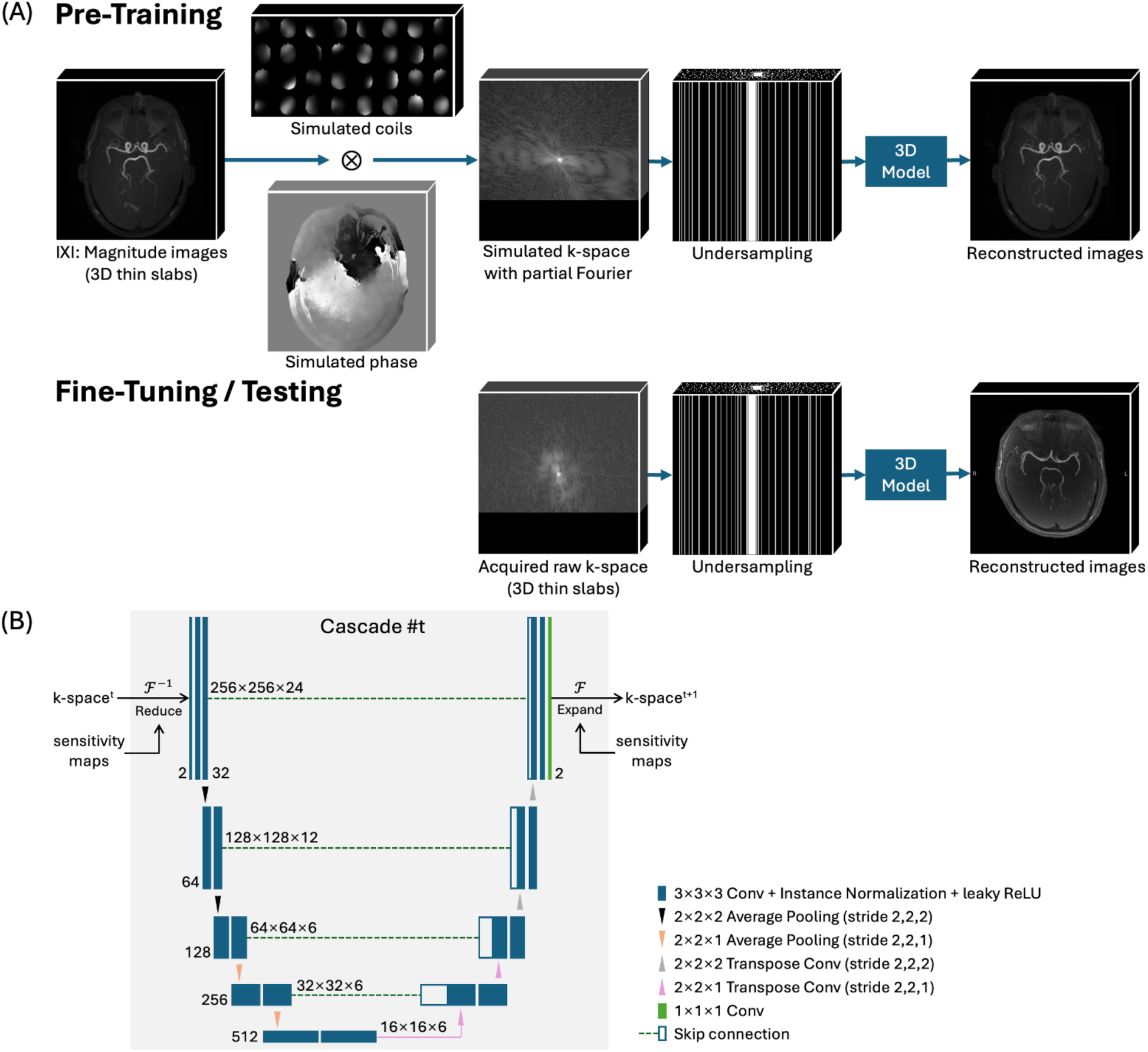
(A) Proposed few-shot learning framework. (B) Proposed model architecture in each cascade in E2E-VarNet. To overcome data limitations, large-scale open-source magnitude images from the IXI dataset were used to simulate complex-valued multi-coil k-space data. Partial Fourier (PF) and Poisson-disc undersampling were then applied. The proposed model integrates 3D pooling and convolution with customized down/up-sampling for thin-slab inputs. After pre-training on simulated k-space data, the model underwent fine-tuning using two single-slab *in vivo* datasets acquired from a 3T scanner. The final model was evaluated on additional experimentally acquired multi-slab *in vivo* k-space datasets.

### 2.1 Raw K-Space Data Simulation

Previous studies have explored various approaches for phase synthesis, including physics-based modeling^37^, sinusoidal functions^38^, and conditional generative adversarial networks (GANs) ^39^. However, GAN-based methods require large raw k-space training datasets, which are generally unavailable for 3D TOF-MRA. Recent efforts have instead leveraged widely available natural images and videos to train DL models for MRI reconstruction, demonstrating promising results. Notably, Wang et al.^40^ simulated the phase by applying Fourier truncation to random complex white Gaussian noise following the LORAKS method^41^. However, their approach borrowed coil sensitivity maps from the 2D fastMRI dataset^22,23,42^ that typically includes at most 16 coil channels with different coverages, which is challenging to adapt to 3D TOF-MRA. Meanwhile, Jaubert et al.^43^ simulated coil sensitivity maps using 2D Gaussians with randomized maximum intensity, standard deviations, and centers. Also, they simulated phase variations relating to specific structures in natural images using their RGB channels, which is not applicable for simulating structure-related phase variations of grayscale MRI images.

In this work, to simulate the phase ϕ realistically, the conjugate symmetric k-space of the magnitude image is corrupted by multiple modulated noise components, resulting in a non-symmetric k-space from which the phase is extracted. The magnitude provides structures to the phase, as observed in experimental data, while the spatially modulated noise components simulate low-frequency background phase variations due to inhomogeneities, and various sources of noise.

The phase simulation model is proposed as:

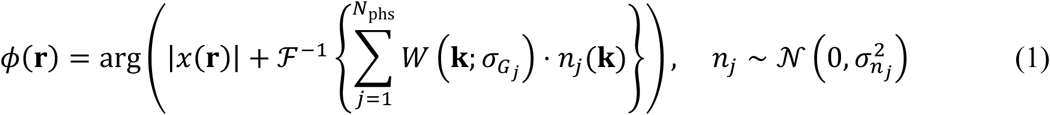

Where:

- ℱ^−1^: The inverse Fourier transform.
- |*x*(**r**)|: The magnitude image with the 3D spatial coordinate **r** = (*r*_x_, *r*_y_, *r*_z_).
- *n*_j_(**k**): A random complex-valued white Gaussian noise at each **k** = (*r*_x_, *r*_y_, *r*_z_) for the *j*-th noise component, sampled from a Gaussian distribution with zero mean and variance 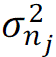.
- *W* (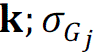): The Gaussian weight map for the *j*-th noise component, centered at (0, 0, 0). *W* is the Gaussian function, where the standard deviation 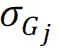 controls the spread of the Gaussian, thereby modulating the spatial frequency of the variations introduced by the noise component.
- *N*_phs_: The total number of distinct noise components added to the phase simulation.

The parameters 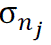 and 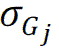 were randomized for each dataset to capture the necessary diversity of variations and were empirically optimized based on reconstruction performance on the validation set, with their specific values and ranges provided in Table S1 of the Supporting Information document. To avoid a phase bias towards zero, the amplitudes of the noise components are set to exceed those of the original signal from the magnitude image.

A representative simulated phase image is shown in Figure 2 alongside the genuine phase from experimental *in vivo* data. The simulated phase demonstrates similar characteristics to the genuine phase, exhibiting slowly varying phase across the head, along with more detailed variations that coincide with vessels and other anatomical structures.

**Figure 2.**
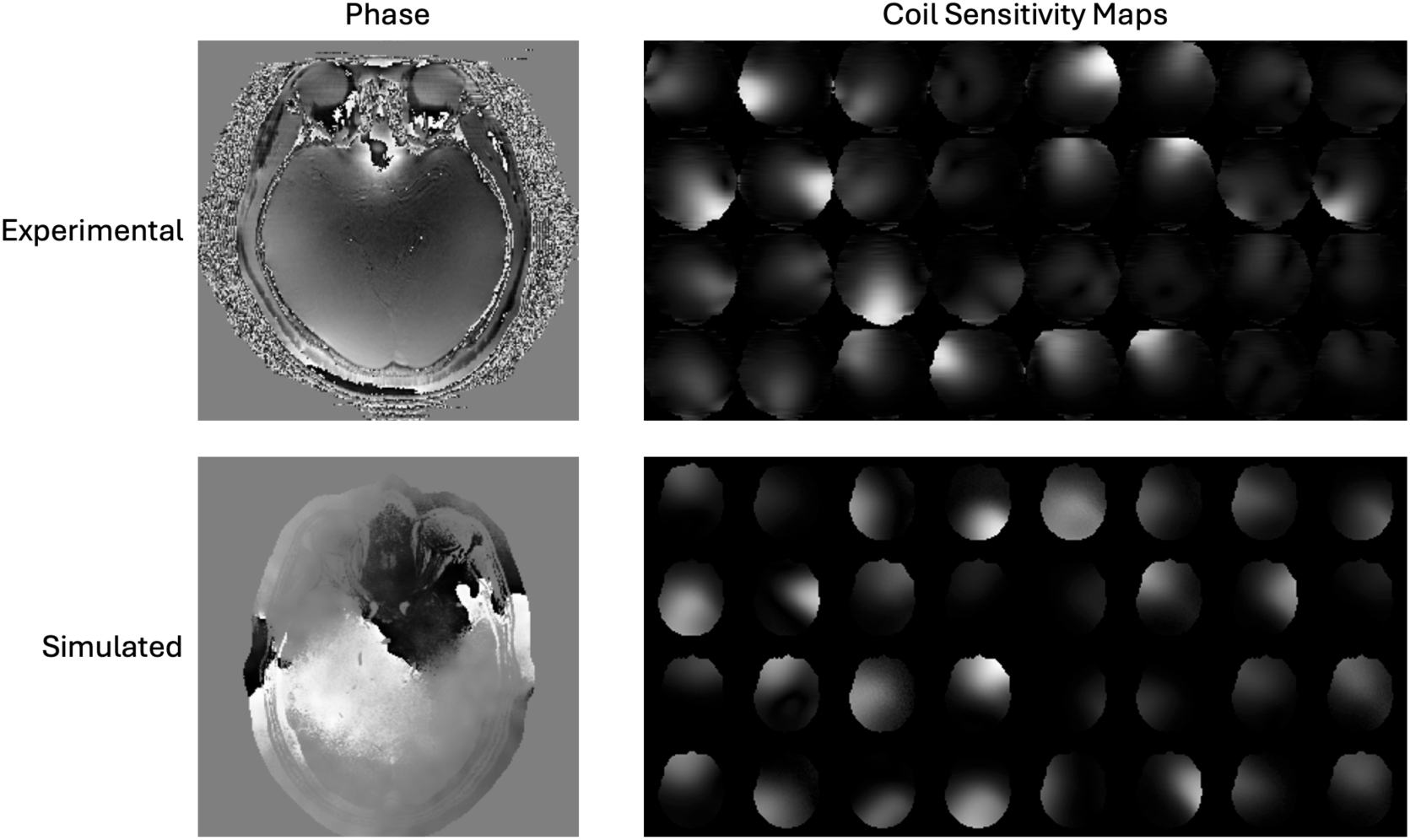
Comparison of simulated phase and coil sensitivity maps with experimentally acquired phase and coil sensitivity maps from *in vivo* data.

To simulate coil sensitivity maps, a head mask *M*_head_(**r**) was first extracted using Otsu’s thresholding followed by morphological operations, including binary erosion with 4 iterations and binary dilation with 9 iterations. The boundary coordinates are then identified using the “find_boundaries” function from the scikit-image library (https://scikit-image.org/). A center location 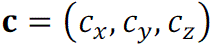 is randomly sampled from the boundaries using a uniform distribution *u*(*boundaries*) for each coil. For the *i*-th coil, a multivariate spatial Gaussian *G*_2_ centered at **c***_i_* is generated and randomly rotated about the z-axis, where 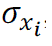, 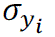, and 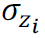 control its spread and shape and are randomly sampled for each coil.

The final sensitivity map for the *i*-th coil is computed as:

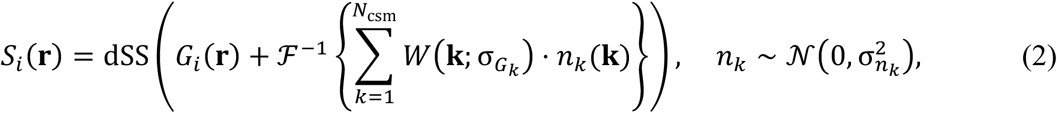

where *N*_csm_ modulated noise components are added to simulate low-frequency background variations and sources of measurement noise, and dSS normalizes the map by the sum-of-squares across coils. Although this formulation is similar to Equation (1), instead of extracting the phase using arg, it simulates the complex-valued low-spatial-frequency coil sensitivity maps directly and adds independent noise to each coil, a critical step as applied in previous work^43^. The Gaussian parameters are listed in Table S2 in the Supporting Information document and representative simulated coil sensitivity maps are shown in Figure 2.

After combining the magnitude image with the simulated phase and 32 coil sensitivity maps, partial Fourier (PF) in the readout direction was simulated, as used in most 3D-TOF-MRA protocols to reduce the TE. To further diversify the dataset, random bias fields caused by field inhomogeneities were introduced using the “RandomBiasField” function (with coefficients = 0.3 and order = 3) from the TorchIO library^44^. Subsequently, 2D pseudorandom variable-density Poisson-disc undersampling masks with a fully sampled calibration region (12 × 6) in the k-space center were applied in the k_y_-k_z_ planes^45^. All simulation processes, excluding the Poisson-disc undersampling, were computed once per dataset and then fixed throughout the training period, while the Poisson-disc undersampling mask was randomized each time a dataset was loaded for training to make the model generalizable to different undersampling patterns.

### 2.2 Model Modifications

To reconstruct 3D TOF-MRA data, where vessel signals exhibit strong continuity across slices, this work extends the physics-informed 2D E2E-VarNet^25^ by implementing 3D pooling and convolution operations throughout all U-Nets. This modification enables the model to effectively capture both intra-and inter-slice correlations, enhancing its ability to reconstruct the complex 3D vasculature in 3D TOF-MRA. Compared to the previous work that used DPI-Net^27^, which utilizes a 3D CNN to learn image-to-image mapping, the proposed model is expected to have improved performance and generalizability by incorporating an MRI forward model.

To further optimize the model for the multiple thin-slab data used in 3D TOF-MRA, its downsampling (max-pooling) and upsampling (up-convolution) layers were carefully designed Specifically, four downsampling levels were used along the longer in-plane axes (x and y), while only two levels were applied along the shorter slice axis to mitigate excessive loss in the through-plane dimension. This architectural modification ensures that sufficient structural information is retained at the bottleneck of the U-Net across the much smaller slice dimension, preventing the degradation of fine vascular details. Additionally, the model enforces PF constraints within each data consistency step, aligning with the readout PF acquisition commonly employed in 3D TOF-MRA protocols.

### 2.3 Training Dataset and *In Vivo* Acquisition

The training dataset was derived from the IXI dataset (https://brain-development.org/ixi-dataset/), comprising images from 341 subjects acquired with a 3T Philips Intera MRI scanner (Philips Medical Systems, Best, The Netherlands), a 1.5T Philips Gyroscan Intera MRI scanner (Philips Medical Systems, Best, The Netherlands), and a 1.5T GE MRI scanner (GE HealthCare, Wisconsin, United States) at Hammersmith Hospital, Guy’s Hospital, and the Institute of Psychiatry in London, United Kingdom. The imaging parameters, as obtained from the dataset’s official website and NIfTI headers, were as follows: repetition time (TR) = 16.7 ms (3T) or 20 ms (1.5T); echo time (TE) = 5.8 ms (3T) or 6.9 ms (1.5T); flip angle = 16° (3T) or 25° (1.5T); acquisition matrix = 288 × 286; voxel resolution = 0.8 × 0.8 × 0.8 mm³; and number of slices = 100. Each 3D magnitude image dataset was divided into five overlapping slabs along the slice axis to replicate multi-slab acquisition, resulting in 24 slices per slab.

To evaluate the proposed DL-based reconstruction method on retrospectively undersampled raw k-space data, five healthy subjects (Cohort 1) were scanned on a 3T Siemens Prisma scanner (Siemens Healthineers, Erlangen, Germany) using a 32-channel receive head coil under a technical development protocol approved by local ethics and institutional committees. The multi-slab 3D TOF-MRA sequence parameters were similar to the IXI data, as follows: TR = 18 ms; TE = 3.8 ms; flip angle = 16°; acquisition matrix = 288 × 288; voxel resolution = 0.8 × 0.8 × 0.8 mm³; and PF = 26% in the readout direction. For each subject, five fully sampled slabs, each consisting of 24 slices with a 20% overlap, were acquired in a total scan time of 12 minutes and 22 seconds.

Additionally, two single-slab datasets were acquired from two other healthy volunteers (Cohort 2) on the same scanner as a validation set for model training. One of these datasets was later used for fine-tuning, while the remaining dataset served as an independent validation set. The single-slab sequence parameters were as follows: TR = 21 ms; TE = 3.48 ms; flip angle = 18°; acquisition matrix = 256 × 256; voxel resolution = 0.8 × 0.8 × 1.0 mm³; PF factor = 29%; number of slices = 20; and acquisition time = 1 minute and 52 seconds.

Finally, prospectively undersampled datasets were acquired using a modified 3D TOF-MRA sequence with predefined pseudorandom variable-density Poisson-disc undersampling masks, using the same imaging parameters as Cohort 1, to demonstrate the real-world potential of this approach. An 8-times accelerated acquisition (scan time 1 min 47 s), along with a fully sampled reference (scan time 12 min 22 s), was obtained from two healthy volunteers (Cohort 3). As the acceleration factor is defined by the undersampling in k-space or, equivalently, the reduction factor in the number of readouts, the slight deviation from an exact 8× reduction in actual scan time results from additional scan overhead, including pre-scans for system calibration.

### 2.4 Experiments

To retrospectively evaluate the reconstruction performance of the proposed method on experimentally acquired *in vivo* 3D TOF-MRA data, quantitative metrics, including the peak signal-to-noise ratio (PSNR)^46^ and structural similarity index measure (SSIM)^47^, were used. These metrics were computed by comparing the entire 3D reconstructed volume with the corresponding fully sampled reference and averaging the results across the dataset. All fully sampled references in this study were obtained by combining the coil images using the conjugate of coil sensitivity maps estimated by ESPIRiT^48^, implemented based on the SigPy package (https://github.com/mikgroup/sigpy).

The proposed method was compared to zero-filling, L1 wavelet-regularized CS^18^, and several DL-based methods, namely: (i) Few-shot DPI-Net (pre-trained and fine-tuned using the proposed few-shot learning framework for 3D TOF-MRA)^27^; (ii) Original 2D E2E-VarNet (originally pre-trained on experimental 2D T1w, T2w, FLAIR raw k-space data from the fastMRI dataset)^25^; (iii) Fine-tuned original 2D E2E-VarNet (the original pre-trained model fine-tuned using two single-slab experimental 3D TOF-MRA datasets); and (iv) Few-shot 2D E2E-VarNet (pre-trained and fine-tuned using the proposed few-shot learning framework for 3D TOF-MRA). For 2D networks, 2D k-space slices were obtained by inverse Fourier transformation along the readout direction. Before image reconstruction, each 32-channel k-space dataset was first cropped to a standard in-plane size of 256 × 256 and compressed into 8 virtual coils to reduce the computational burden using an SVD-based method^49^.

All models were pre-trained for 50 epochs for each undersampling factor (4× and 8×) using Adam optimizers, a 0.0003 learning rate, validation-based early stopping, and L1+SSIM loss minimization on an NVIDIA A100 80GB GPU. The proposed 3D model took approximately 3.5 days to pre-train, while 2D models took around 1.5 days. Fine-tuning on an individual experimental single-slab dataset from the validation set (Cohort 2) followed the same protocol as pre-training (50 epochs, learning rate = 0.0003, validation-based early stopping, and L1+SSIM loss). Fine-tuning required approximately 10 minutes for 3D models and 8 minutes for 2D models.

To further assess the proposed method, ablation experiments were conducted. First, the proposed 3D model was trained using only two single-slab experimental 3D TOF-MRA datasets—one for fine-tuning and one for validation—without any pre-training on simulated raw k-space data (named Proposed w/o pre-training) to evaluate the importance of pre-training. Second, the proposed model was pre-trained on simulated data but was not fine-tuned (named Proposed w/o fine-tuning) to assess the impact of fine-tuning.

To evaluate the impact of the proposed raw k-space data simulation approach, the proposed 3D model was pre-trained on two alternative datasets generated using recent methods. Specifically, these two datasets replaced part of the proposed simulation pipeline with the phase simulation method from LORAKS^41^, as recently used by Wang et al.^40^, and the recent coil sensitivity map simulation method from Jaubert et al.^43^, respectively.

Finally, to assess the performance and robustness of the proposed method on prospectively undersampled data, reconstructed images were visually compared with their fully sampled references. Quantitative metrics were not used in this case due to the potential for subject motion, physiological changes, and scanner drift between the prospectively undersampled and fully sampled acquisitions.

## 3 Results

### 3.1 Comparison of Retrospective Dataset Reconstructions

Two acceleration factors, R=4 and R=8, were tested retrospectively on the experimentally acquired *in vivo* k-space data, with representative axial maximum intensity projection (MIP) reconstructions shown in Figure 3 and Figure 4. Reconstruction times for each multi-slab dataset on an NVIDIA H100 10GB GPU were approximately 125 seconds for compressed sensing, 12 seconds for Few-shot DPI-Net, and 15 seconds for both 2D E2E-VarNets and the proposed method.

**Figure 3.**
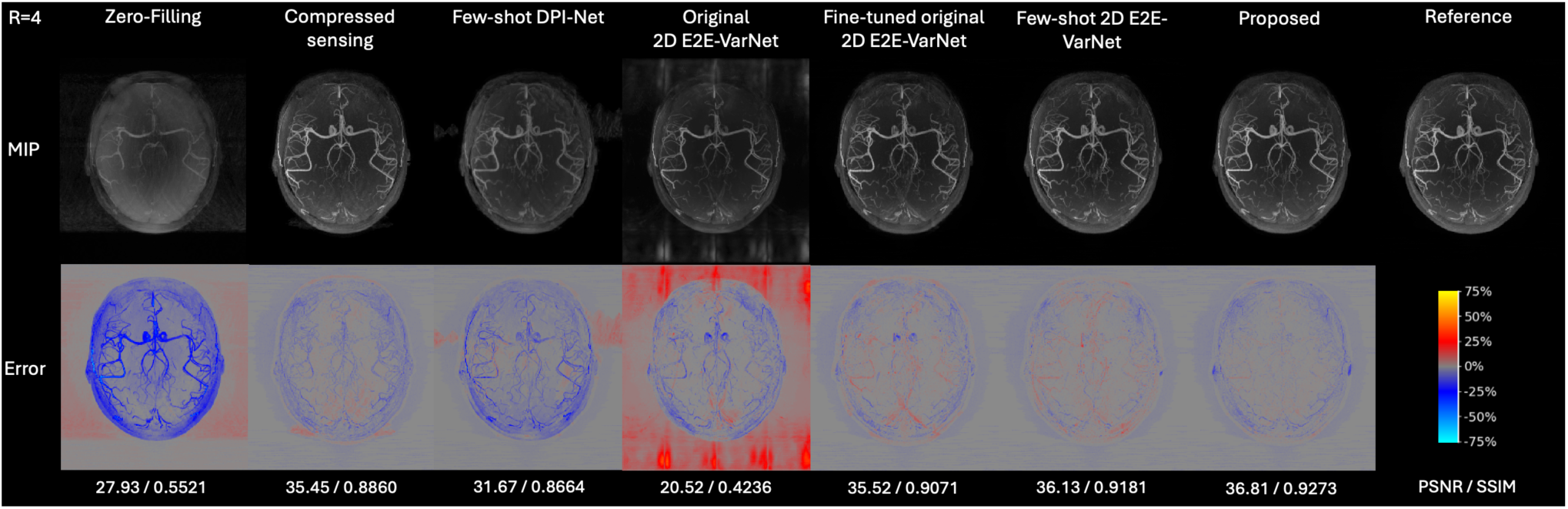
Axial MIP reconstructions of retrospectively undersampled experimental *in vivo* data for acceleration factor R=4, showing results for comparison methods alongside the fully sampled reference. The top row shows reconstructed MIP angiograms (greyscale-adjusted for comparison), the second row shows error maps (percentage difference from the reference normalized by the maximum intensity), and PSNR/SSIM metrics are reported at the bottom.

**Figure 4.**
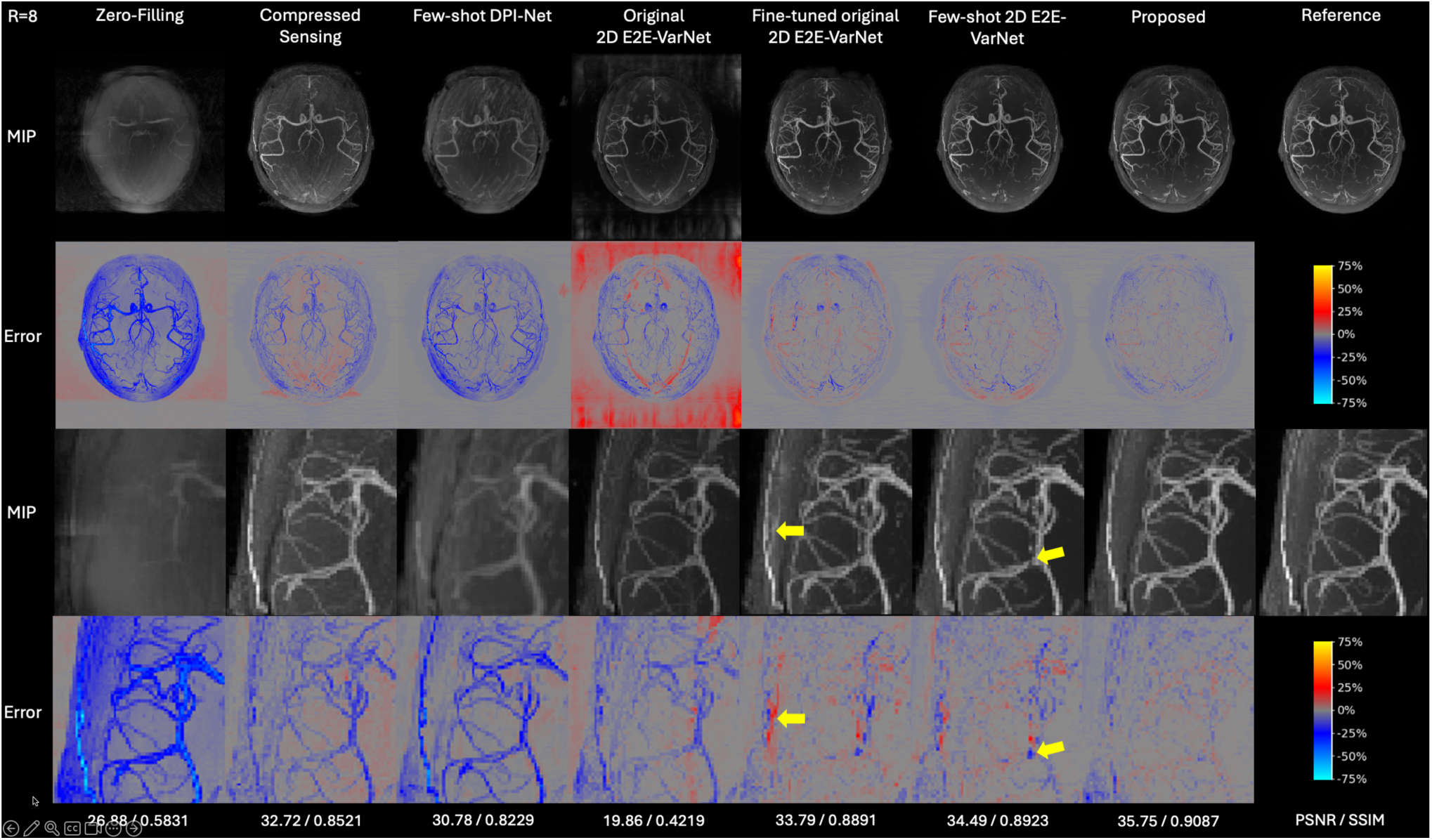
Axial MIP reconstructions of retrospectively undersampled experimental *in vivo* data for R=8 with zoomed views. The first two rows follow Figure 3, the third and fourth rows are zoomed views of the first two rows, and PSNR/SSIM metrics are shown at the bottom. Yellow arrows indicate artifacts in Fine-tuned original 2D E2E-VarNet and missed vessel signals in Few-shot 2D E2E-VarNet as examples.

The proposed method demonstrated superior image quality, retaining most fine vessels with minimal noise and artifacts, especially for R=8. In contrast, other methods exhibited higher noise and aliasing, particularly Original 2D E2E-VarNet and Fine-tuned original 2D E2E-VarNet. However, pre-training on the simulated 3D TOF-MRA raw k-space dataset via the proposed few-shot learning framework significantly enhanced the performance of 2D E2E-VarNet, though some aliasing artifacts still remained. Few-shot DPI-Net exhibited a relatively large degradation in reconstruction quality from R=4 to R=8, with noticeable blurring and missing vessels. Quantitative results in Table 1 further highlight the greater improvements achieved by the proposed method with R=8, where it improved PSNR by 1.77dB and SSIM by 2.66% compared to the Fine-tuned original 2D E2E-VarNet.

**Table 1.**
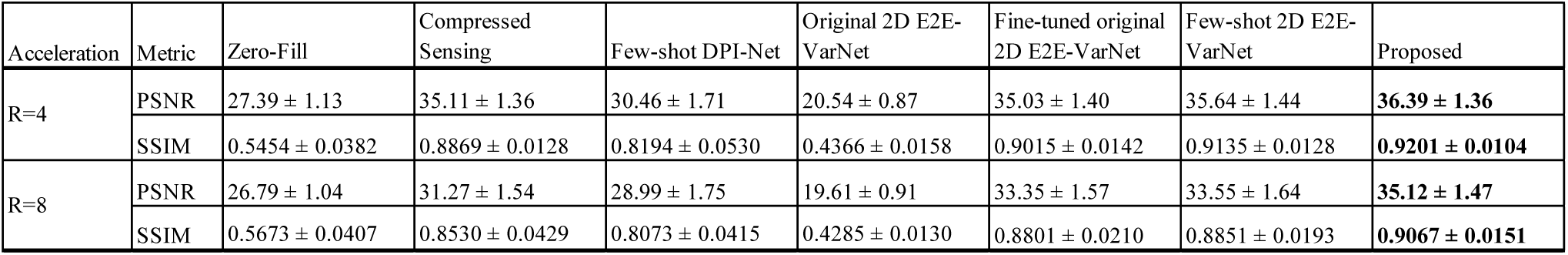
Quantitative comparison of reconstruction performance on retrospectively undersampled experimental *in vivo* data for R=4 and R=8, showing results for methods in Figure 3 and Figure 4. The table shows PSNR and SSIM metrics, presented as mean ± standard deviation across all subjects. The top-performing method for each task and metric is highlighted in bold font.

### 3.2 Ablation Experiments

To separately assess the benefits of pre-training on extensive simulated data and fine-tuning on a single slab of experimental *in vivo* data, representative MIP reconstructions and quantitative results for methods including Proposed w/o pre-training, Proposed w/o fine-tuning, and Proposed are shown in Figure 5 and Table 2. Proposed w/o pre-training exhibited significantly lower reconstruction quality for both R=4 and R=8 accelerations, highlighting the importance of the proposed pre-training approach. While Proposed w/o fine-tuning achieved good reconstruction performance, fine-tuning using just two single-slab experimental datasets further improved reconstruction for both acceleration factors, effectively reducing noise and yielding higher PSNR and SSIM.

**Figure 5.**
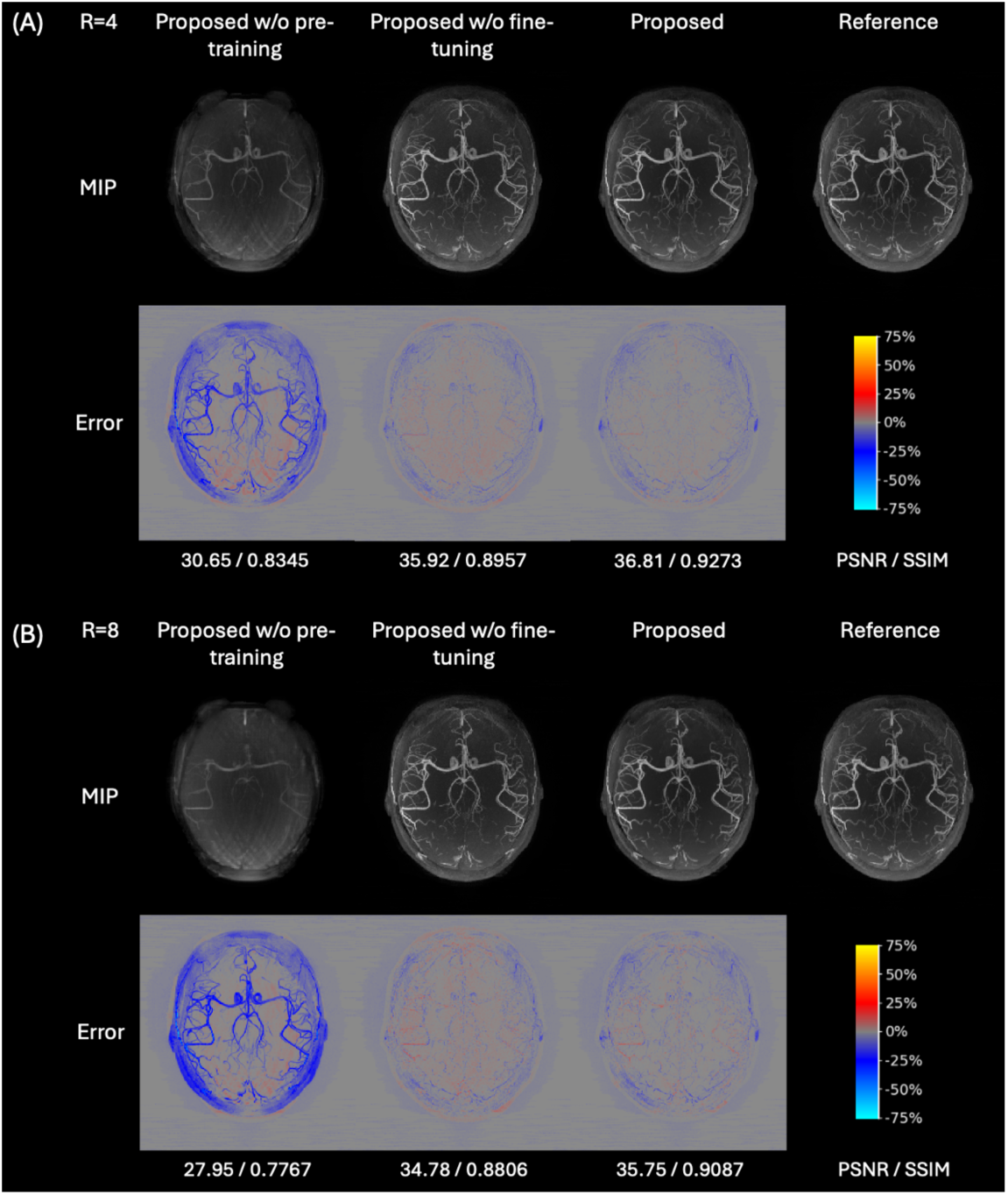
Axial MIP reconstructions of retrospectively undersampled experimental *in vivo* data for R=4 (subfigure A) and R=8 (subfigure B), showing results for methods: Proposed w/o pre-training, Proposed w/o fine-tuning, and Proposed. The subfigure layout follows Figure 3.

**Table 2.**
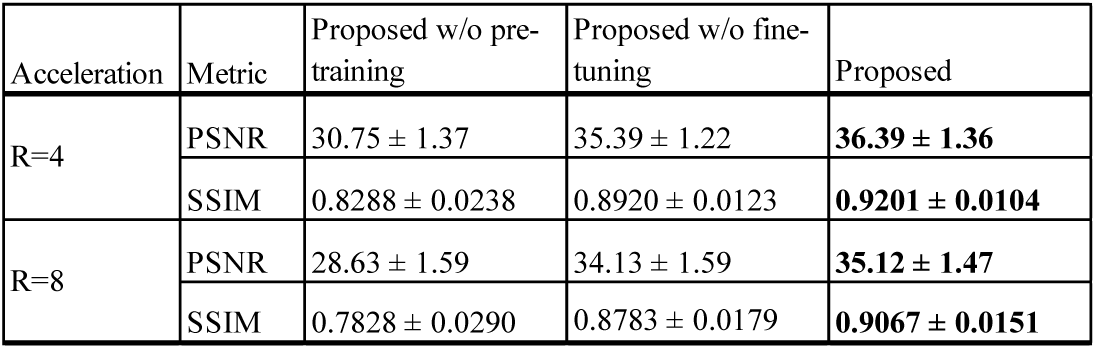
Quantitative comparison of reconstruction performance on retrospectively undersampled experimental *in vivo* data for R=4 and R=8, showing results of methods in Figure 5. The table format follows Table 1.

### 3.3 Comparison of Simulation Methods

To evaluate the impact of the proposed raw k-space data simulation methods, MIP reconstructions and quantitative results of the proposed 3D model, trained on two alternative datasets that replaced part of the simulation pipeline with two adapted recent methods (LORAKS^41^ and Jaubert et al.^43^), are compared in Figure 6 and Table 3 for R=8. The proposed data simulation pipeline demonstrated higher reconstruction quality compared to existing simulation methods, achieving the highest PSNR and SSIM.

**Figure 6.**
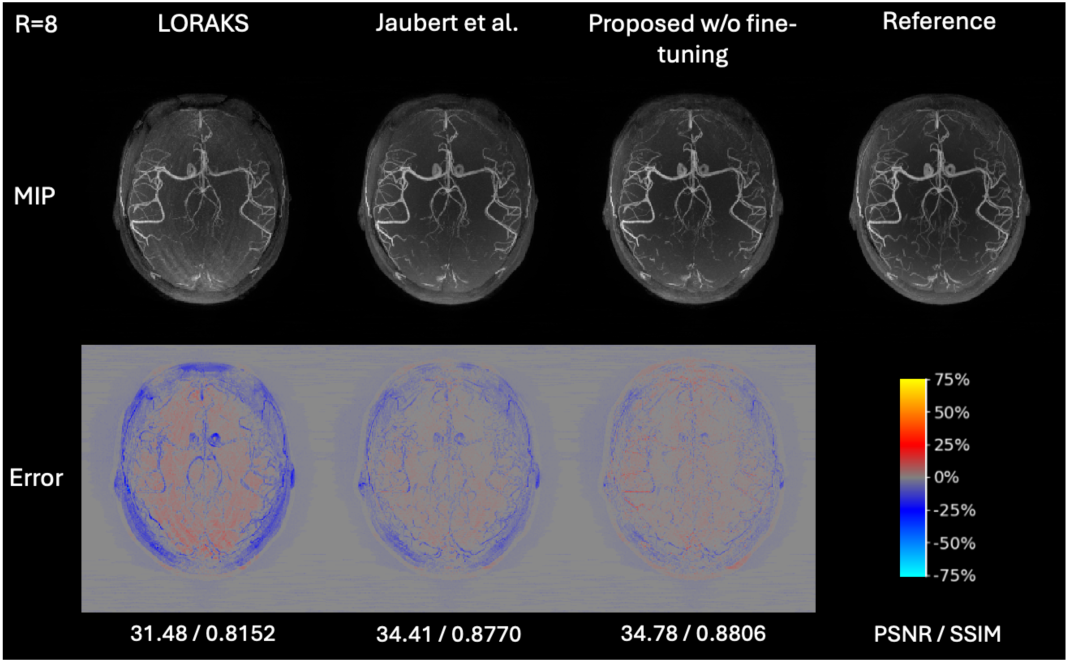
Axial MIP reconstructions of retrospectively undersampled experimental *in vivo* data for R=8, showing results of the proposed 3D model trained on the dataset generated by the proposed data simulation pipeline and datasets where part of the pipeline was replaced with methods from LORAKS and Jaubert et al. The figure layout follows Figure 3.

**Table 3.** Quantitative comparison of reconstruction performance on retrospectively undersampled experimental *in vivo* data for R=8, showing results of methods in Figure 6. The table format follows Table 1.

| Acceleration | Metric | LORAKS | Jaubert et al. | Proposed w/o fine-tuning |
| --- | --- | --- | --- | --- |
| R=8 | PSNR | $30.76 \pm 1.74$ | $33.73 \pm 1.47$ | **$34.13 \pm 1.59$** |
| | SSIM | $0.8088 \pm 0.0327$ | $0.8762 \pm 0.0171$ | **$0.8783 \pm 0.0179$** |

### 3.4 Reconstructions of Prospective Datasets

To evaluate the performance and robustness of the proposed methods in a real-world scenario, the reconstruction results of the Proposed and Proposed w/o fine-tuning methods on the prospectively 8-fold accelerated dataset were compared against Zero-Filling, L1 wavelet-regularized Compressed Sensing, and fully sampled references, as shown in Figure 7. Compared to Compressed Sensing, both proposed methods reconstructed more vessels with fewer artifacts and aliasing, consistent with the previous retrospective studies. Notably, the Proposed, fine-tuned using only two single slabs of experimental data, demonstrated the highest apparent signal-to-noise ratio (SNR). As highlighted by the yellow arrows in Figure 7, single-slab fine-tuning reduced noise and made the small vessel signals more clearly visible. However, some very small vessels were still lost, and slight residual aliasing persisted in the posterior region of Subject 2, as indicated by the red arrows in Figure 7. This may be attributed to motion-related data quality degradation, as lower SNR was also observed in Compressed Sensing reconstructions for Subject 2.

**Figure 7.**
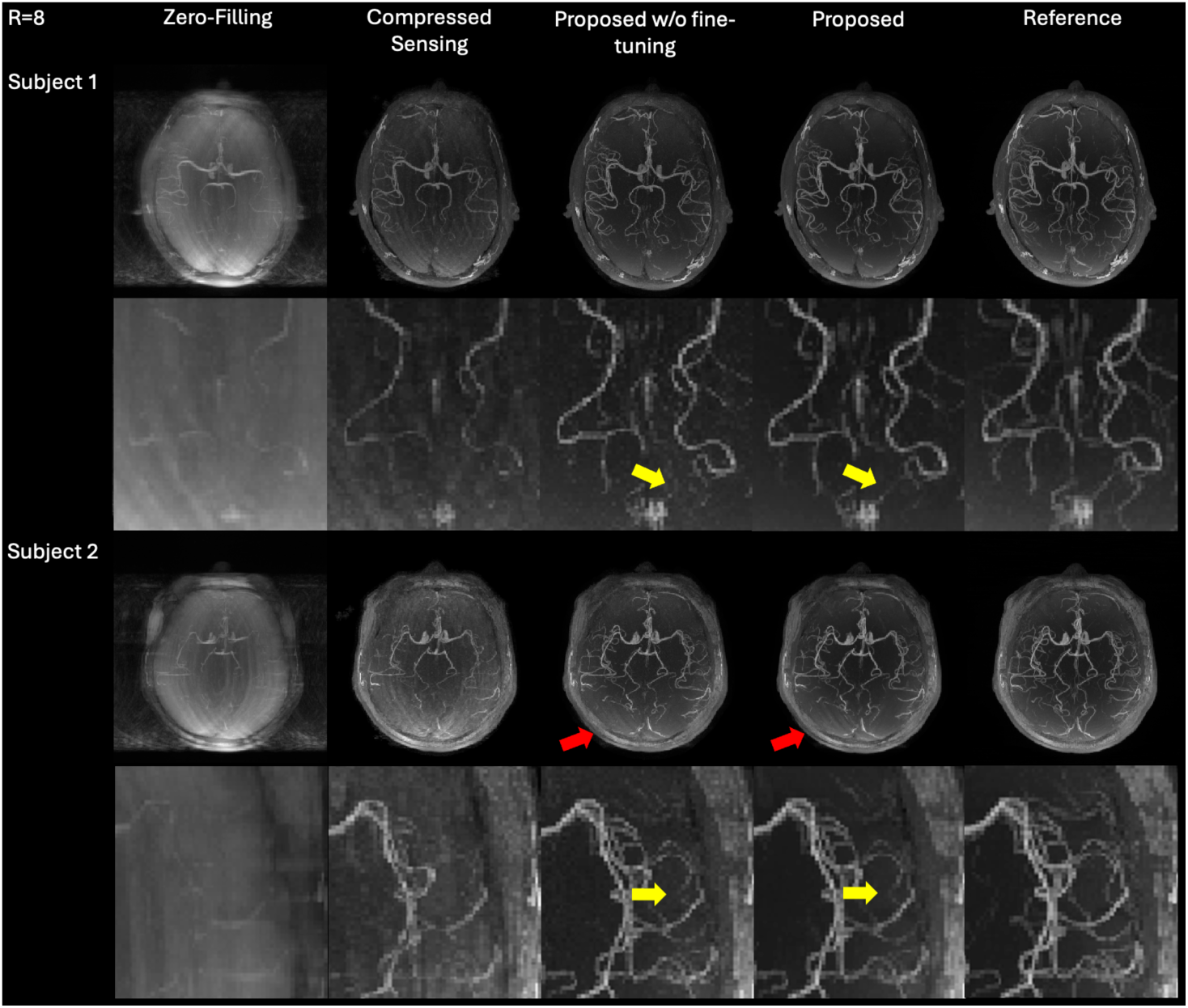
Axial MIP reconstructions of prospectively undersampled experimental *in vivo* data for R=8 with zoomed views, showing results for Zero-Filling, L1 wavelet-regularized Compressed Sensing, Proposed w/o fine-tuning, and Proposed methods. For each subject, the first row presents reconstructed MIP angiograms, and the second row provides zoomed views. Yellow arrows indicate improvements due to single-slab fine-tuning, while red arrows highlight slight residual aliasing.

## 4 Discussion

The proposed few-shot learning approach for highly accelerated 3D TOF-MRA, integrating a novel raw k-space data simulation method with an adapted 3D variational network model, effectively reconstructed high-quality 3D TOF-MRA images from undersampled *in vivo* acquisitions relying on only two single-slab experimentally acquired raw k-space datasets for fine-tuning and validation. It maintained high image fidelity even under more aggressive undersampling (8x), with rapid reconstruction times (15s for all slabs).

In retrospective studies, the proposed method outperformed all other tested reconstruction methods in reconstructing retrospectively undersampled experimental *in vivo* 3D TOF-MRA data, particularly at higher acceleration factors, which exhibited minimal noise and artifacts with only obvious missing details in non-critical peripheral regions such as the ears and skull. In contrast, the 2D E2E-VarNet pre-trained solely on the fastMRI dataset with T1w, T2w, and FLAIR images exhibited persistent aliasing even after fine-tuning. However, the 2D E2E-VarNet pre-trained on simulated 3D TOF-MRA raw k-space datasets achieved much less aliasing and higher image quality, which underscored the value of the proposed few-shot learning framework for robust generalization. However, this advantage did not extend to DPI-Net, which may be attributed to its direct image-to-image mappings without incorporating MRI physics models, making it more susceptible to overfitting.

The effectiveness of the proposed 3D model, incorporating architectural modifications tailored to 3D TOF-MRA, was evident when compared to the 2D E2E-VarNet. The customized 3D model leveraged both inter– and intra-slice correlations for continuous vessel depiction and was specifically designed to better handle 3D thin-slab inputs with partial Fourier, underscoring the importance of modality-specific model adaptations.

Ablation experiments demonstrated that the proposed pre-trained model already achieved strong performance, and fine-tuning using just two single-slab experimental *in vivo* datasets further improved reconstruction quality by effectively reducing aliasing and noise. This highlights the potential generalization ability of the proposed approach to new scanners and protocols, where two single-slab datasets could be used to adapt the pre-trained model to new conditions, avoiding the need for large training datasets to be acquired at each site. Additionally, the experiments underscored the value of the proposed few-shot learning framework in data-limited scenarios, as training an effective reconstruction model from scratch using only very limited experimentally acquired data proved infeasible.

Compared to other raw k-space data simulation techniques, the proposed simulation method enabled the model to reconstruct 3D TOF-MRA images with higher quality and fewer artifacts. This demonstrates the ability of the proposed data simulation pipeline to generate more realistic raw k-space data for 3D TOF-MRA through its more complex and curated simulation processes, which account for high-frequency variations inherent to its flow-related enhancement nature. The proposed pipeline demonstrated a more significant improvement over existing methods in phase simulation, which is inherently more challenging than coil sensitivity simulation due to the absence of smooth and consistent profiles across MRI modalities. Therefore, enhancing phase simulation is likely to be more critical, and the new method introduced here, with broad parameter ranges, effectively captures the necessary diversity in phase profiles.

When tested on prospectively undersampled datasets, the proposed method consistently demonstrated high-quality reconstructions. Compared to the benchmark method, Compressed Sensing, the proposed method reconstructed more fine vessels with markedly less noise and aliasing. The robust and consistent performance of the proposed method on retrospective studies and prospectively undersampled datasets—similar to real-world scenarios—can be attributed to the use of large amounts of multi-center, multi-vendor training magnitude images and the proposed randomized data simulation with wide parameter ranges. Additionally, the benefit of fine-tuning the proposed method using only two single-slab experimental *in vivo* datasets was also seen in prospectively undersampled data.

Despite these strengths, some limitations remain. First, for the acceleration factor of 8, although the proposed method significantly outperformed other methods, some very small vessel signals were missed in the reconstructions, particularly for the method without fine-tuning. This may be due to a residual domain shift between the simulated and experimental acquired data, which could potentially be addressed by a more rigorous optimization of all parameters within the data simulation pipeline or by incorporating additional data sources for better generalization. Second, when data quality is low— such as in cases of poor signal-to-noise ratio or significant motion corruption—some aliasing remains in the reconstructions. This issue could be mitigated by incorporating simulations of motion corruption and employing other data augmentation techniques in future work. Third, the proposed methods were evaluated on a limited healthy cohort, and further validation is needed with a larger, more diverse patient population, particularly those with various cerebrovascular disorders, to better assess its clinical utility. Finally, developing an inline reconstruction pipeline would be essential in future work to enhance the clinical applicability of the proposed methods.

In conclusion, this work has demonstrated the effectiveness of a new few-shot learning-based reconstruction method designed for highly accelerated 3D TOF-MRA, which requires only minimal experimentally acquired data to achieve superior results on both retrospective and prospective *in vivo* data over existing methods with rapid reconstruction times. Given the challenges of collecting a large set of raw k-space data for 3D TOF-MRA, this method holds significant promise for advancing research and clinical applications of high-resolution, whole-head 3D TOF-MRA imaging.

## Acknowledgments

The Wellcome Centre for Integrative Neuroimaging is supported by core funding from the Wellcome Trust (203139/Z/16/Z and 203139/A/16/Z) with additional support from the NIHR Oxford Biomedical Research Centre (NIHR203311) and the Oxford Health Biomedical Research Centre (NIHR203316). H.L. is supported by the Medical Research Council, the Nuffield Department of Clinical Neurosciences, and Siemens Healthineers through an Oxford-MRC iCASE studentship.

M.C. is supported by the Canada Research Chairs Program. P.J. is supported by the Dunhill Medical Trust and the NIHR Oxford Biomedical Research Centre. T.O. is supported by a Sir Henry Dale Fellowship jointly funded by the Wellcome Trust and the Royal Society (220204/Z/20/Z).

## Conflict of Interest

Hao Li receives studentship support from Siemens Healthineers. Iulius Dragonu is an employee of Siemens Healthineers. Peter Jezzard is the Editor-in-Chief of Magnetic Resonance in Medicine. In line with COPE guidelines, Peter Jezzard recused himself from all involvement in the review process of this paper, which was handled by an associate editor. He and the other authors had no access to the identities of the reviewers.

## Data Availability Statement

The original IXI dataset used for training can be found here: https://brain-development.org/ixi-dataset/. The code underlying the phase and coil simulations and model training, as well as the pre-trained model, can be found here <LINK to be added upon publication>. We are currently unable to share the full in vivo data due to data protection issues, although the Wellcome Centre for Integrative Neuroimaging is actively working on a solution to this.

