## Supporting Information for "Few-shot learning for highly accelerated 3D time-of-flight MRA reconstruction"

Table S1. Parameters for phase simulation. Values for $\sigma_{n_j}$ represent ratios relative to the maximum intensity, while values for $\sigma_{G_j}$ represent ratios relative to the size of the dimensions. $\mathcal{U}(a, b)$ denotes a uniform distribution within a range $[a, b]$ from which the parameters are randomly sampled.

| $j$ | $\sigma_{n_j}$ | $\sigma_{G_j}$ |
|---|---|---|
| 1 | $\mathcal{U}$(1.0, 3.0) | $\mathcal{U}$(0.005, 0.025) |
| 2 | $\mathcal{U}$(0, 0.01) | $\mathcal{U}$(0.025, 0.25) |
| 3 | $\mathcal{U}$(0, 0.0001) | 1.0 |

Table S2. Parameters for coil sensitivity map simulation. Values for $\sigma_{n_j}$ represent ratios relative to the maximum intensity, while values for $\sigma_{G_j}$, $\sigma_{x_i}$, $\sigma_{y_i}$, and $\sigma_{z_i}$ represent ratios relative to the size of the dimensions. $\mathcal{U}(a, b)$ denotes a uniform distribution within a range $[a, b]$ from which the parameters are randomly sampled, while $\mathcal{N}(\mu, \sigma)$ is a Gaussian distribution with mean $\mu$ and standard deviation $\sigma$.

| $j$ | $\sigma_{n_j}$ | $\sigma_{G_j}$ | $\sigma_{x_i}$ | $\sigma_{y_i}$ | $\sigma_{z_i}$ |
|---|---|---|---|---|---|
| 1 | $\mathcal{U}$(0, 0.25) | 0.0075 | $\mathcal{N}$(0.25, 0.05) | $\mathcal{N}$(0.25, 0.05) | $\mathcal{N}$(0.25, 0.05) |
| 2 | $\mathcal{U}$(0, 0.0001) | 1.0 | | | |
